# Historical isolates provide insights into the evolution of *Acinetobacter baumannii* international clone 2 and its resistome

**DOI:** 10.64898/2026.01.20.700532

**Authors:** Matthew Neil, Frédéric Grenier, Nancy Allard, Gemma C. Langridge, Santiago Castillo-Ramírez, Louis-Patrick Haraoui, Benjamin A. Evans

**Author notes:** Corresponding author. Mailing address: Norwich Medical School, University of East Anglia, Bob Champion Research and Education Building (BCRE), Rosalind Franklin Road, Norwich Research Park, Norwich, NR4 7UQ, UK.

## Abstract

*Acinetobacter baumannii* remains a pervasive nosocomial pathogen, with international clone 2 (IC2) being the most successful lineage. IC2 is often discussed as a singular entity, and the genomic factors behind its success over other lineages are poorly characterised. This study aimed to track genomic changes occurring around the emergence of IC2 and further determine the within-clone population structure. A set of historical *A. baumannii* isolates from the 1970s to early 2000s were long-read sequenced and assembled into high-quality chromosomes. These were then used to supplement publicly available *A. baumannii* genomes, producing a dataset of 1,281 chromosomes derived from isolates collected across six continents. A recombination-free phylogeny was produced and combined with antimicrobial resistance gene (ARG) presence/absence data. Analysis showed that IC2 began to emerge as the dominant lineage around 2005, which coincides with the acquisition of *oxa23* and AbGRI3. Further, IC2 can be split into at least four distinct groups. Groups 1, 2, and 3 represent incremental evolution of the *A. baumannii* chromosome over time, while group 4 represents a separate evolutionary path distinct from the main IC2 lineage, which has recently been isolated at higher frequency.

**Significance statement:** Characterising the evolution and population structure of antimicrobial-resistant bacterial pathogens is of key importance both for epidemiological monitoring and developing approaches to combat infections and their spread. Here, we use a unique collection of historical isolates to show that the major international clone in the WHO priority pathogen *Acinetobacter baumannii* is comprised of different, independent groups with individual antimicrobial resistance gene compliments. This highlights new areas for research to understand the clinical impact of the different groups and the drivers underlying the evolution and spread of epidemic clones of antimicrobial-resistance bacterial pathogens.

## Introduction

Antimicrobial resistance (AMR) remains one of the most substantial causes of human morbidity and mortality worldwide, with *Acinetobacter baumannii* being particularly notable for its AMR properties. In 2019 alone, it was responsible for >400,000 AMR-related deaths, making it the fifth most lethal bacterial pathogen by this metric (1). In addition to being a pervasive nosocomial pathogen, *A. baumannii* can be found in animals, plants, food products and even aquatic environments, making it a One Health issue (2). Despite its current ubiquity, *A. baumannii* is thought to have emerged relatively recently, likely around the 1970s (3,4). During this early expansion, international clone 1 (IC1) and IC2 lineages were the dominant form of *A. baumannii*. Then, IC2 began to overtake IC1 as the dominant lineage, and currently accounts for the majority of *A. baumannii* infections worldwide, followed by IC5 (5). However, the lack of sequencing data from historical isolates means that identifying the genomic changes underlying the success of IC2 over IC1 is nearly impossible to determine with existing data.

The *A. baumannii* genome is characterised by a small core genome and a large, diverse accessory genome containing an array of antimicrobial resistance genes (ARGs) and virulence factors (VFs) that tend to be concentrated on the chromosome (6–8). Intrinsic ARGs in *A. baumannii* encode class C and class D β-lactamases, referred to as *Acinetobacter*-derived cephalosporinase (ADC) and *Acinetobacter baumannii* oxacillinase (OxaAb, also referred to as OXA-51-like) enzymes, respectively. Additionally, the *A. baumannii* chromosome contains clusters of genes involved in the biosynthesis of extracellular capsular polysaccharides (CPS) and lipooligosaccharides (LOS), known as the K-locus and OC-locus. These play a major role in virulence and resistance to both environmental stresses and antibiotics (9,10). *A. baumannii* chromosomes also contain a vast array of acquired ARGs. Two of the most clinically relevant are *oxa23* (also referred to as *bla_oxa-23_*), conferring high-level carbapenem resistance (11), and *armA*, which confers resistance to a broad range of aminoglycosides (12,13). Due to the high genome plasticity of *A. baumannii*, closely related isolates may have very different ARGs, leading to potentially different antimicrobial susceptibility patterns.

Many ARGs in *A. baumannii* are co-localised in resistance islands (RIs), of which there are two main types: AbaR-type RIs are associated with IC1, while AbGRI-type RIs are associated with IC2 (14). Of the AbGRI-type RIs, AbGRI3 is perhaps the most clinically relevant due to the presence of *armA*, which is found exclusively within AbGRI3 in IC2 isolates (15). AbGRI3 contains three main sections: a core containing *armA*, *mphE*, and *msrE*; a large integron containing *sul1*, *qacEΔ1*, *aadA*, *catB8*, and *acc(6*’*)-Ib9* (also referred to as *aacA4*), which together with the core region forms Tn*6180*; and an additional transposon (Tn*6179*) containing *aph(3*’*)-Ia* (also referred to as *aphA1b*). At least five variants of AbGRI3 exist based on the presence/absence of these sections and the deletion of a replication initiation protein in the core section and/or a class 1 integron integrase in the integron section (15). However, for simplicity, here we define “short” AbGRI3 variants as those which only contain the core section (version 4 in Blackwell *et al*. (15)), and “long” AbGRI3 variants as those which contain additional sections.

Within the core genome, *A. baumannii* displays low genetic variance due to high levels of recombination and recent, rapid clonal expansion (16). This low variance makes finding subgroups within IC2 challenging. Nonetheless, an approach using recombination-free alignments has previously been applied to IC1 to track its emergence and the genomic factors behind its success (17). A recent paper by Li *et al*. (18) similarly found subgroups within IC2. However, they did not fully characterise the genome content of each group with respect to AMR, even though isolates separated by little evolutionary distance in the core genome can have considerable variation in their accessory genome, which includes ARGs (19). Therefore, it is unclear which ARGs may be associated with each group, and what potential variation between insertion sequence (IS)/RI content exists between groups.

To gain a better understanding of the emergence and structure of IC2 over time, we sequenced a set of historical *A. baumannii* genomes. These were combined with publicly available sequences to produce a comprehensive dataset of *A. baumannii* chromosomes collected across six continents and five decades. By integrating ARG presence/absence with a recombination-free phylogeny, we show that IC2 can be split into at least four distinct groups. Three of these groups reflect a stepwise acquisition and loss of certain ARGs over time, associated with changes in the RI AbGRI3. The fourth group is distinct from the other three, representing a parallel evolutionary path within the *A. baumannii* IC2 major lineage.

## Results

### Historical *A. baumannii* chromosomes suggest the emergence of IC2 began in 2005

To analyse the changes in the *A. baumannii* chromosome over time, a set of 226 historical *A. baumannii* isolates from 1970s-early 2000s were collated. All isolates were sequenced using Oxford Nanopore Technology and assembled into complete chromosomes. All chromosomes showed benchmarking universal single-copy orthologs (BUSCO) scores >95% and average nucleotide identity (ANI) >95% when compared to the reference strain ATCC 19606. This set of isolates was then combined with publicly available *A. baumannii* genomes at “complete” or “chromosome” level to create a dataset of 1,281 chromosomes. The majority of isolates were collected from the late 1970s to the present day, although 7 chromosomes were from isolates collected before 1970. Of the 148 isolates sequenced in this study with a known country of origin, most were from Europe (48%) and the United Kingdom (UK) (17%), with smaller contributions from countries such as Turkey (11%), Singapore (6%), and Argentina (6%). Meanwhile, the majority of isolates collected after 2010 were from the USA and China, reflecting global trends in strain isolation and genome sequencing.

All isolates were assigned to ICs using multi-locus sequence typing (MLST) according to the Pasteur scheme. As expected, IC2 was the largest, with just over half (50.9%, 628 isolates) of all isolates assigned a sequence type (ST) belonging to IC2. During inspection of the core genome phylogeny, it was found that one IC2 isolate was not grouped with other IC2 isolates and formed a distinct outgroup, so it was removed from downstream analyses. The first IC2 isolate in this dataset was collected in 1982. However, it was not until the early 2000s (∼2005) that IC2 overtook IC1 to become the dominant clone (Figure 1). Since then, IC2 has shown rapid growth, while IC1 has shown far less growth. Therefore, it is probable that any genomic changes which underlie IC2’s success would have arisen around the early 2000’s.

**Figure 1.**
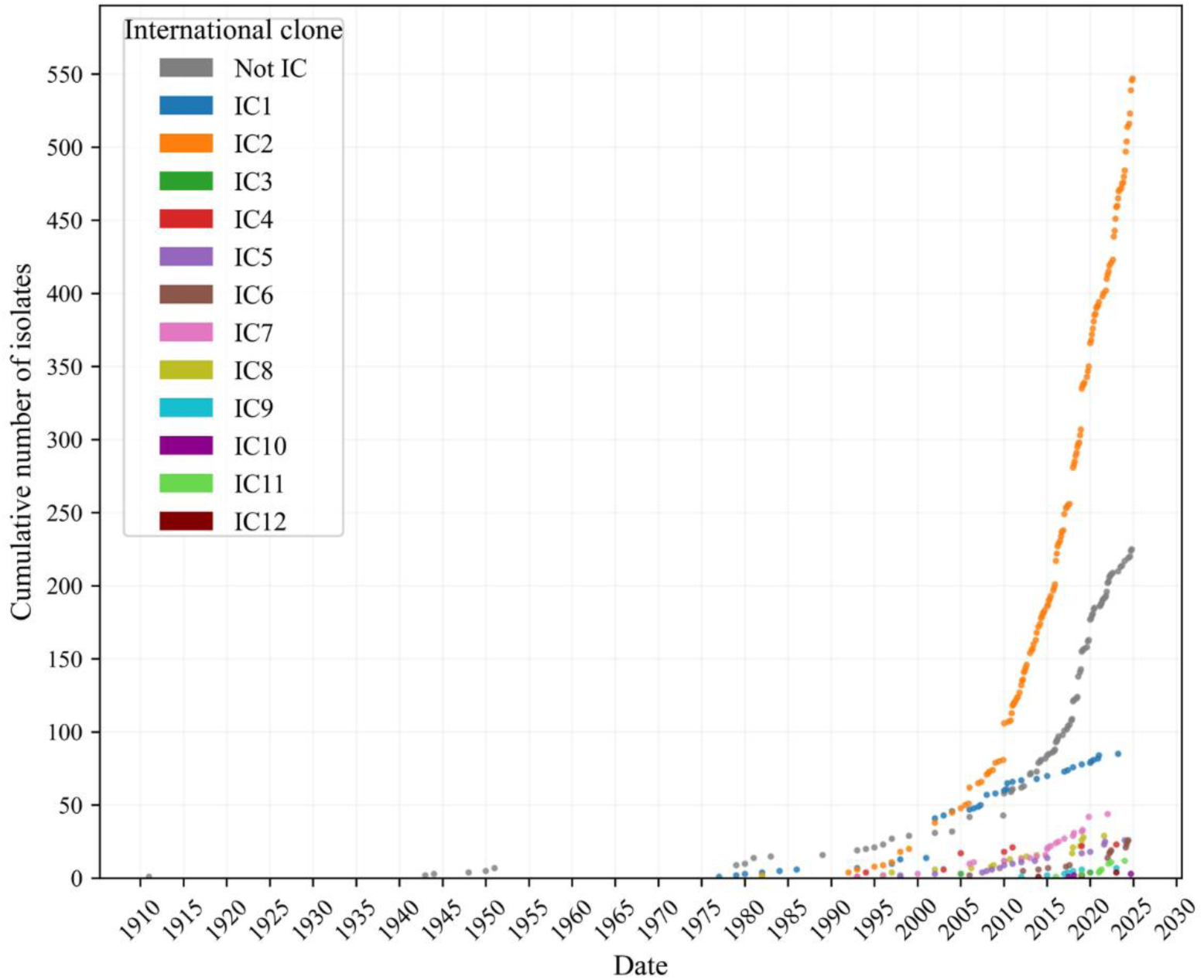
Cumulative number of isolates over time. Each dot represents a single isolate. Dots are colored according to assigned IC, with isolates not belonging to any IC shown in grey.

### IC2 isolates are enriched in a diverse set of ARGs

To further investigate the genomic changes in IC2 that may have contributed to its success, ARGs were annotated using the resistance gene identifier (RGI), which queries the comprehensive antibiotic resistance database (CARD). The highest number of ARGs per chromosome was found in a set of three IC1 isolates with a large gene amplification (20) (GenBank accession no. CP090607.1, CP090606.1, CP091172.1). Beyond these three isolates, IC2 isolates were found to have a significantly higher number of ARGs on their chromosomes than all other ICs and those not belonging to an IC (Figure 2). Further, a large set of ARGs was significantly enriched in IC2 (Figure 3), many of which were either unique to this clone or were only sparsely found outside of it. These ARGs confer resistance to a wide array of antimicrobials, including carbapenems, aminoglycosides, sulphonamides, fluoroquinolones, macrolides, and tetracyclines, as well as certain disinfecting agents. This, combined with the higher median number of chromosomally encoded ARGs in IC2 isolates over other IC/not IC isolates, suggests that IC2’s success may have been driven by resistance to a variety of antimicrobials rather than to a single antimicrobial.

**Figure 2.**
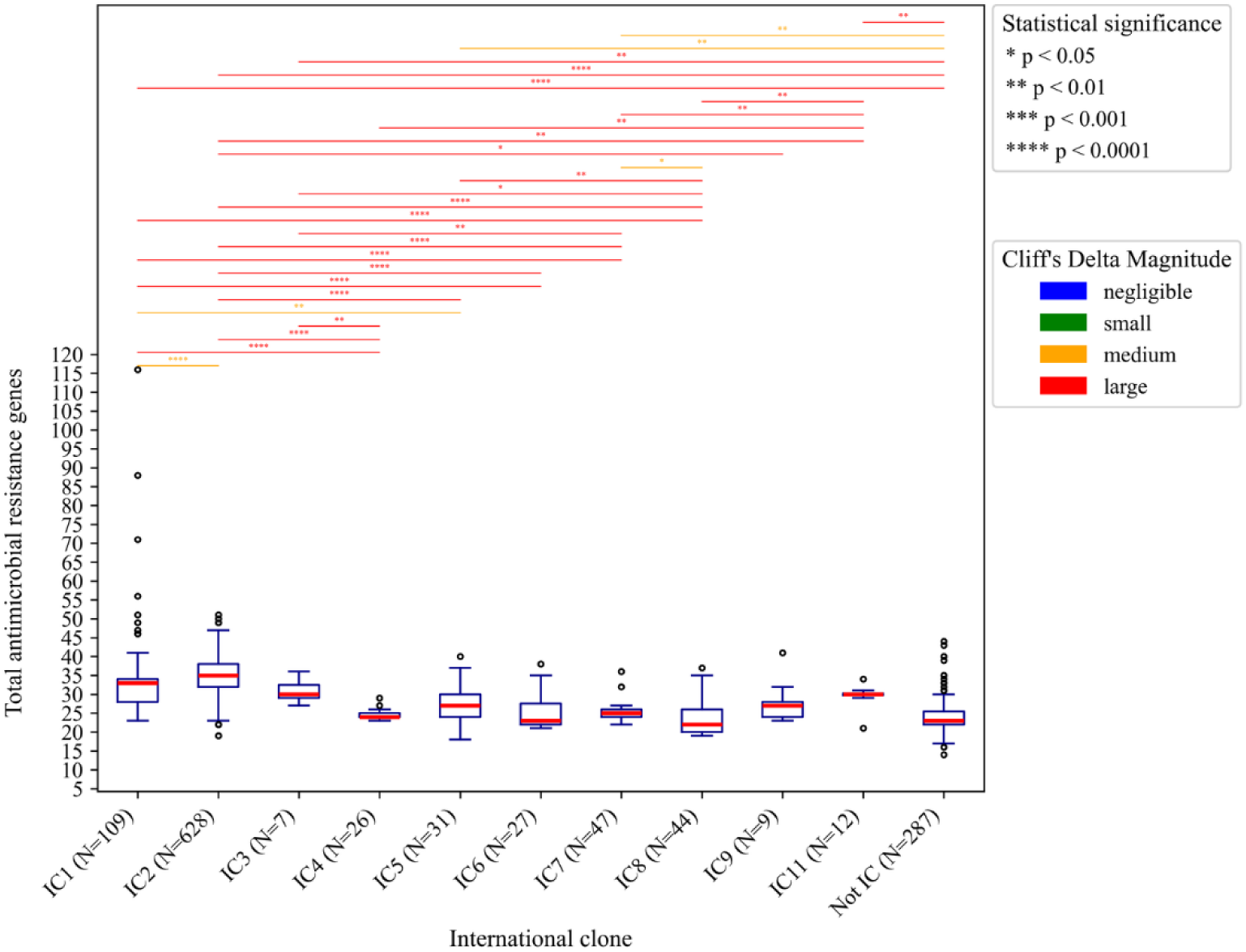
Boxplots of total ARGs on *A. baumannii* chromosomes in the different ICs. IC10 and IC12 been omitted due to being represented by fewer than 5 isolates in our dataset. Medians are shown with red lines. Outliers (±1.5×IQR) are shown as open circles. Statistical significance (*p*<0.05) was determined using the Mann-Whitney U test with the Holm-Bonferroni correction applied.. Cliff’s delta was used to determine the effect size.

**Figure 3.**
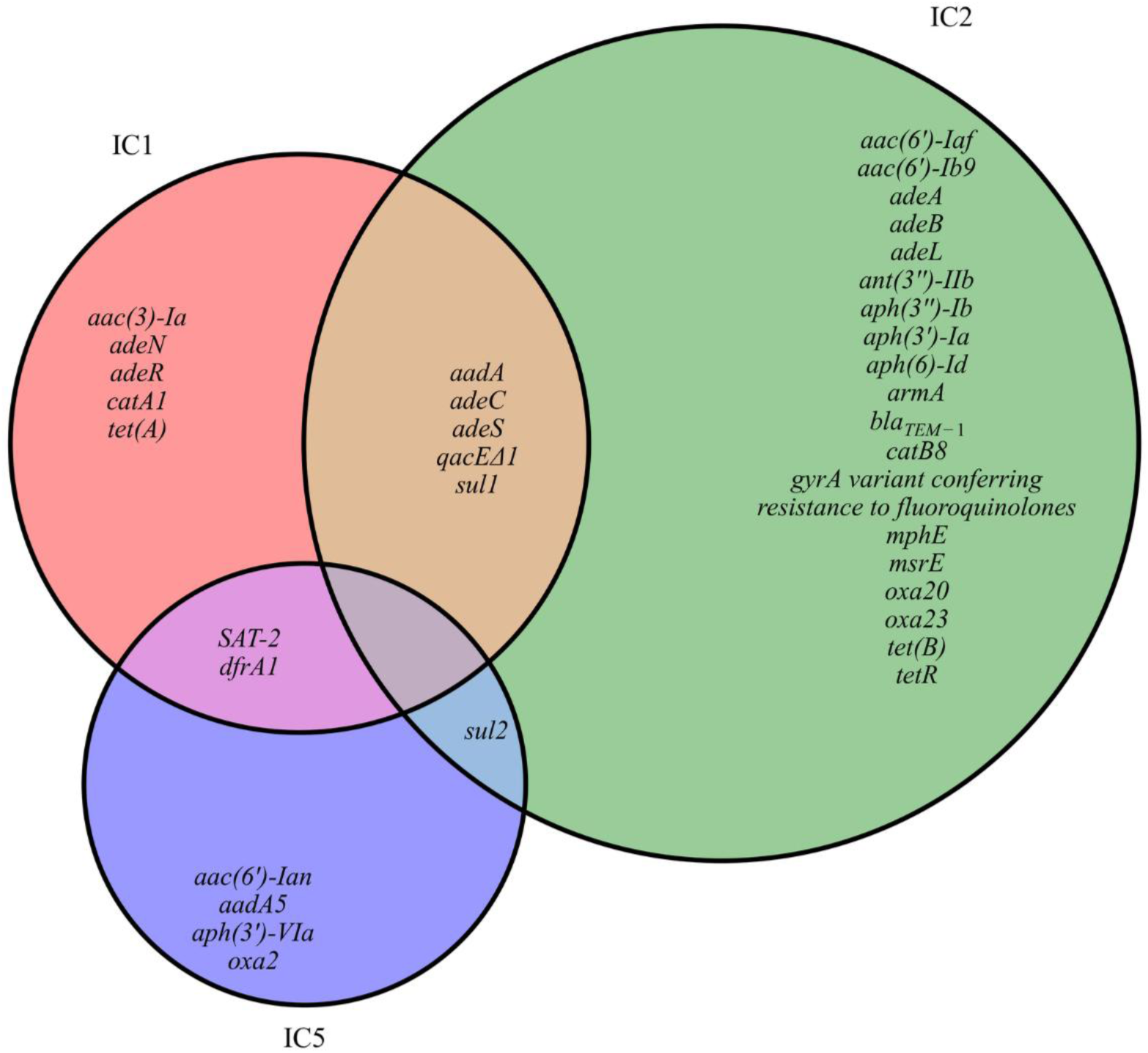
Enrichment of ARGs in IC1, IC2, and IC5. ARGs were annotated using RGI. Statistically significant (*p*<0.05) enrichment of ARGs in each IC was determined using Fischer’s exact test with the Holm-Bonferroni correction applied. Results from the three most globally dominant ICs (IC1, IC2, and IC5) are shown, with other ICs omitted. The intrinsic *bla*_ADC_ and *oxaAb* genes have been omitted.

### IC2 can be split into at least four groups with corresponding ARG presence/absence patterns

Following the previous analysis of ARG enrichment in IC2, the proportions of IC2 isolates containing ARGs over time were investigated. Two patterns were noted amongst these ARGs. A subset of ARGs (*armA*, *mphE*, *msrE*, and *oxa23*) are nearly all absent from IC2 until around 2005, after which the proportion of isolates containing these genes increases rapidly, with the genes remaining common in IC2 to this day (Figure 4A). A different set of genes (*catB8*, *aac(6*’*)-Ib9*, *bla_ADC-30_*, *aadA*, *sul1*, and *qacEΔ1*), some of which also show rapid acquisition after around 2005, drop considerably in proportion after about 2015 and do not recover (Figure 4B). These patterns of ARG presence/absence show that the genome content of IC2 has been changing dramatically over time. Further, the fact that the acquisition of some of these genes coincides with the sudden expansion of IC2 suggests that this subset of ARGs may have contributed significantly to the success of IC2.

**Figure 4.**
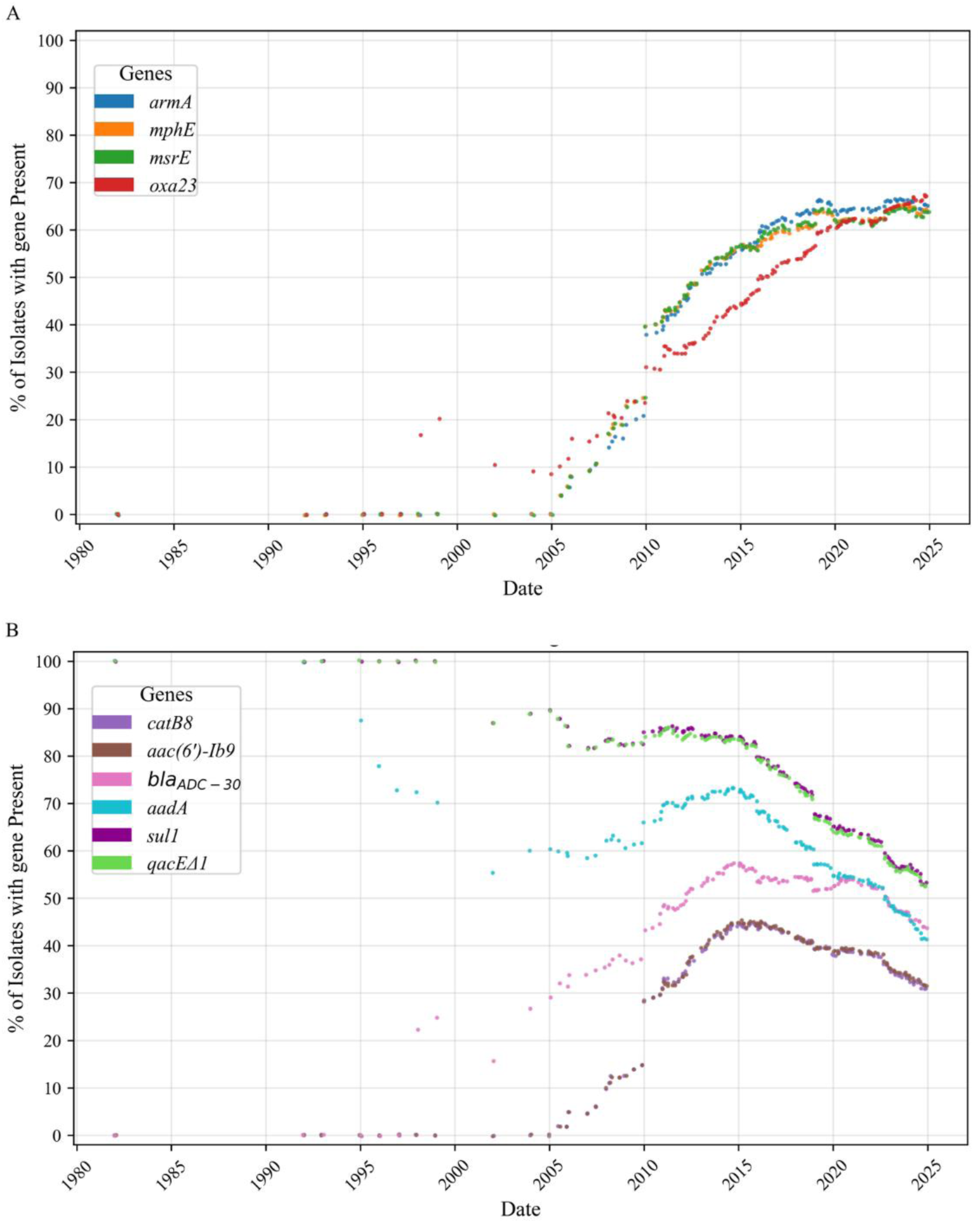
Proportion of IC2 isolates containing ARGs over time. Each dot represents the percentage of IC2 isolates (N=627) up to and including that date which contain at least one chromosomal copy of the corresponding ARG. (A) and (B) show different sets of ARGs with differing presence/absence patterns over time. In (A), percentage values plateau towards a maximum value due to the inclusion of older isolates which lack the ARGs shown. Dots are colored according to a corresponding ARG.

To further investigate potential groups within IC2, an IC2 phylogeny was created from a recombination-free alignment and rooted with ATCC 19606 as an outgroup (see Methods). Root-to-tip analysis gave an R^2^ value of 0.14, which was insufficient for dating. However, bootstrap support values were high across the phylogeny, especially in shallow branches, giving high confidence in major splits. Examining this phylogeny revealed a number of clades with shared ARG presence/absence patterns, splitting IC2 into at least four groups (Figure 5). The occurrence of each group over time is shown in Figure 6. Group 1 contains the oldest isolates and contains few IC2-specific ARGs aside from *aadA*, *sul1*, and *qacEΔ1*, all of which are also enriched in IC1 (Figure 3), suggesting these genes may be ancestral to both IC2 and IC1. Group 2 contains the next most recent isolates, being the dominant IC2 group between 2005-2015, and accounting for a high proportion of IC2 isolates to this day. Most of its isolates contain the ARGs discussed previously (*armA*, *mphE*, *msrE*, and *oxa23*), accounting for their sudden increase in frequency around 2005 (Figure 4). Group 3 represents the most recent clade of IC2 to emerge. While the first isolate was found in 2005, the group did not start rapidly expanding until approximately 2015. Of the ARGs specifically enriched in IC2, only the subset *armA*, *mphE*, *msrE*, and *oxa23* are widespread in these isolates, which accounts for the steep decline in frequency of other ARGs. The final major group, Group 4, appears to have emerged independently of groups 2 and 3. While the first isolate was collected in 2007, the phylogeny suggests it is the oldest. There has been a notable increase in the isolation of *A. baumannii* belonging to group 4 since 2022. However, these recent isolates originated from the Utah Public Health Laboratory Infectious Disease submission group or the Centers for Disease Control and Prevention, both in the USA. Therefore, it is unclear whether this represents a recent expansion (as seen in groups 2 and 3 in 2005 and 2015, respectively), or is due to increased pathogen surveillance. Aside from the ARGs described above, group 4 isolates are deficient in almost all ARGs which are enriched in IC2 over other ICs. This is with the notable exception of *oxa23*, which is common in isolates from this group.

**Figure 5.**
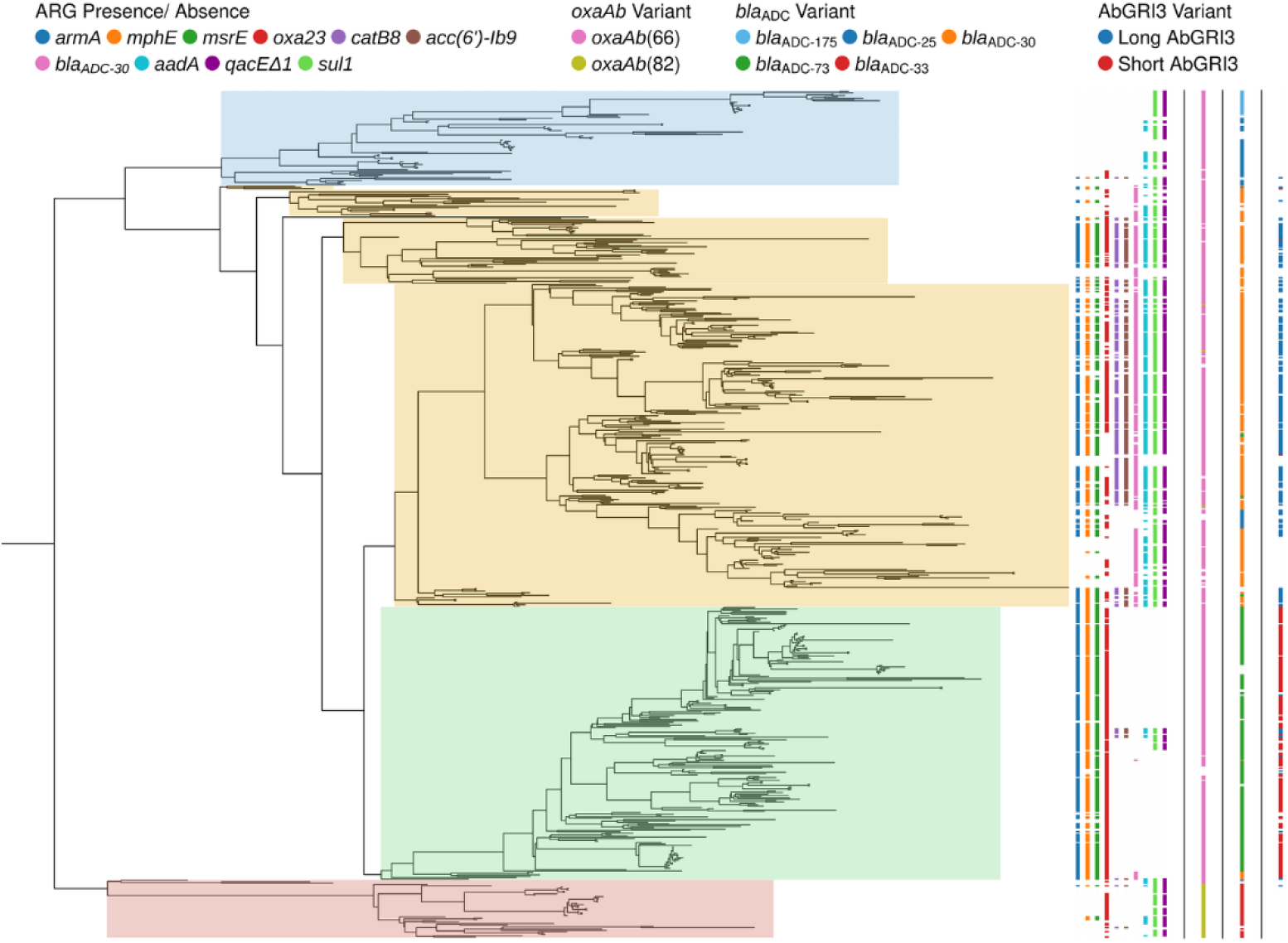
IC2 phylogeny highlighting groups. The phylogeny was created using Gubbins and outgroup rooted as described in Methods. The four major groups are marked by background shading of the phylogeny: group 1 in blue, group 2 in yellow, group 3 in green, and group 4 in red. The four datasets to the right of the phylogeny represent, from left to right, the presence of ARGs, the *oxaAb* variant, the *bla*_ADC_ variant, and the AbGRI3 variant. For simplicity, only “short” and “long” AbGRI3 variants were identified as stated in the Introduction. Only major *oxaAb* and *bla*_ADC_ variants are shown.

**Figure 6.**
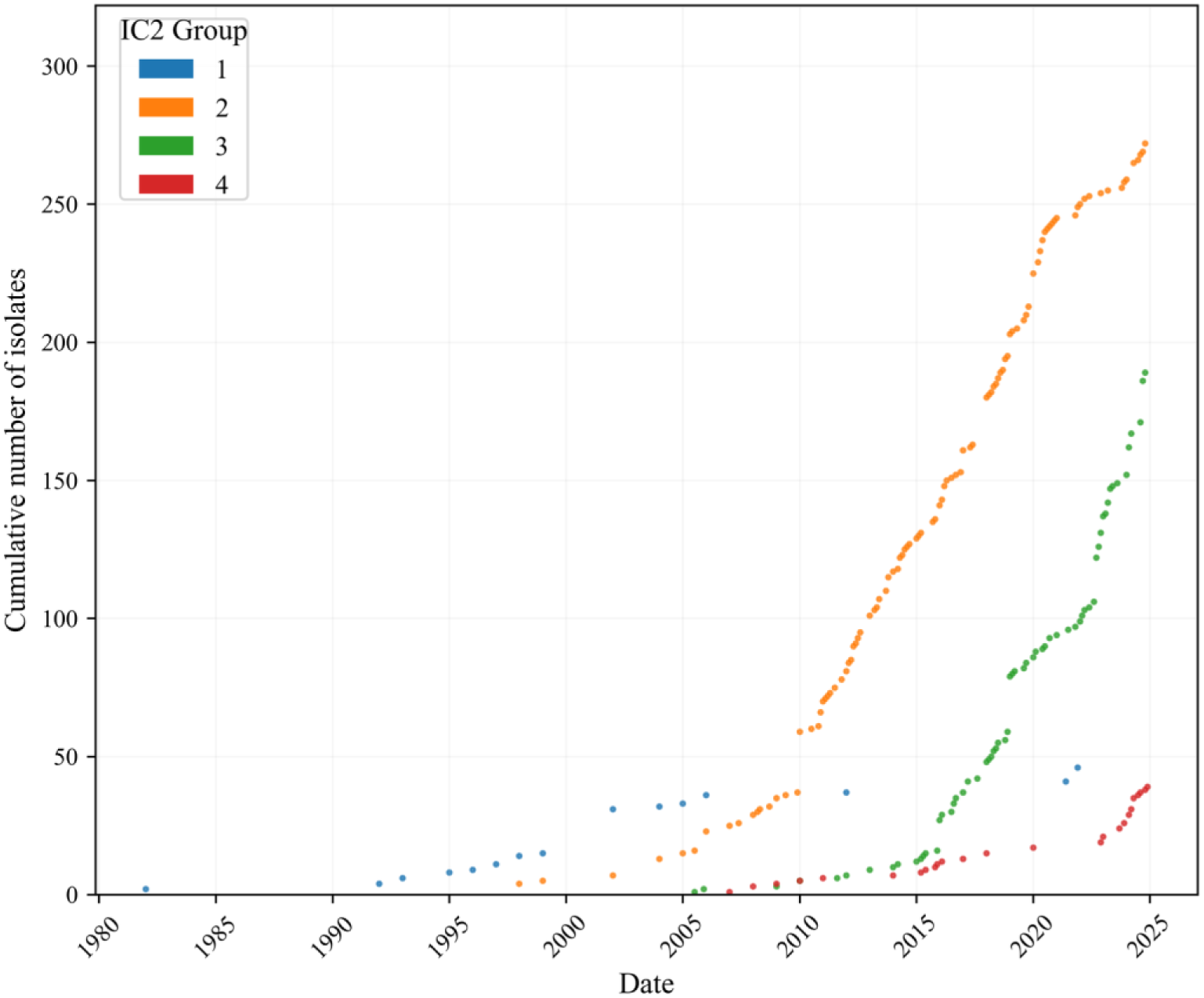
Cumulative number of IC2 group isolates over time. Each dot represents a single isolate and is colored according to the groups shown in the legend.

### IC2 groups contain distinct variants of AbGRI3

Many of the ARGs (*acc(6*’*)-Ib9*, *catB8*, *aadA*, *qacEΔ1*, *sul1*, *armA*, *msrE*, *mphE*) that show group-specific presence/absence patterns are contained within AbGRI3. Due to the significant variance in the content and exact structure of AbGRI3 between individual isolates, we identified only “short” and “long” AbGRI3 variants as described in the Introduction. Groups 2 and 3 were enriched in long and short variants, respectively (Figure 5). Since the AbGRI3-localised genes which are absent from group 3 (*acc(6*’*)-Ib9*, *catB8*, *aadA*, *qacEΔ1*, *sul1*) are found in a single integron, the switch from long to short AbGRI3 likely occurred in a single deletion event.

### *oxaAb* and *bla*_ADC_ gene distribution shows group 4 is distinct from other IC2 groups

While ARG presence/absence patterns show a clear distinction between IC2 groups, which is congruent with the IC2 population structure represented by the recombination-free phylogeny, further analysis was performed to confirm how these groups are related by analysing the *oxaAb* and *bla*_ADC_ gene variants. The *bla*_ADC_ genes showed greater variation: group 1 contained mostly *bla_ADC-25_*, with a separate subgroup containing *bla*_ADC-175_, groups 2 and 3 contained *bla_ADC-30_* and *bla_ADC-73_*, respectively, and group 4 mainly contained *bla_ADC-33_* (Figure 5). Analysis of amino acid differences between these genes suggests an evolution from *bla_ADC-25_* to *bla_ADC-30_* to *bla_ADC-73_*, with each gene differing from its predecessor by only a single amino acid substitution (Figure 7A). *bla_ADC-33_* was most similar to *bla_ADC-25_* with only 2 amino acid substitutions (Figure 7A). *bla_ADC-175_* was also most similar to *bla_ADC-25_*, but by 8 substitutions which were distributed across the gene (Figure 7B), suggesting significant evolutionary distance between these isolates and the rest of group 1. Overall, this is consistent with the inference from the data on group prevalence (Figure 6), AMR gene content, and AbGRI3 structures (Figure 5) that groups 1, 2, and 3 arose sequentially. Further, it supports the phylogeny in suggesting that group 4 is distinct from other groups.

**Figure 7.**
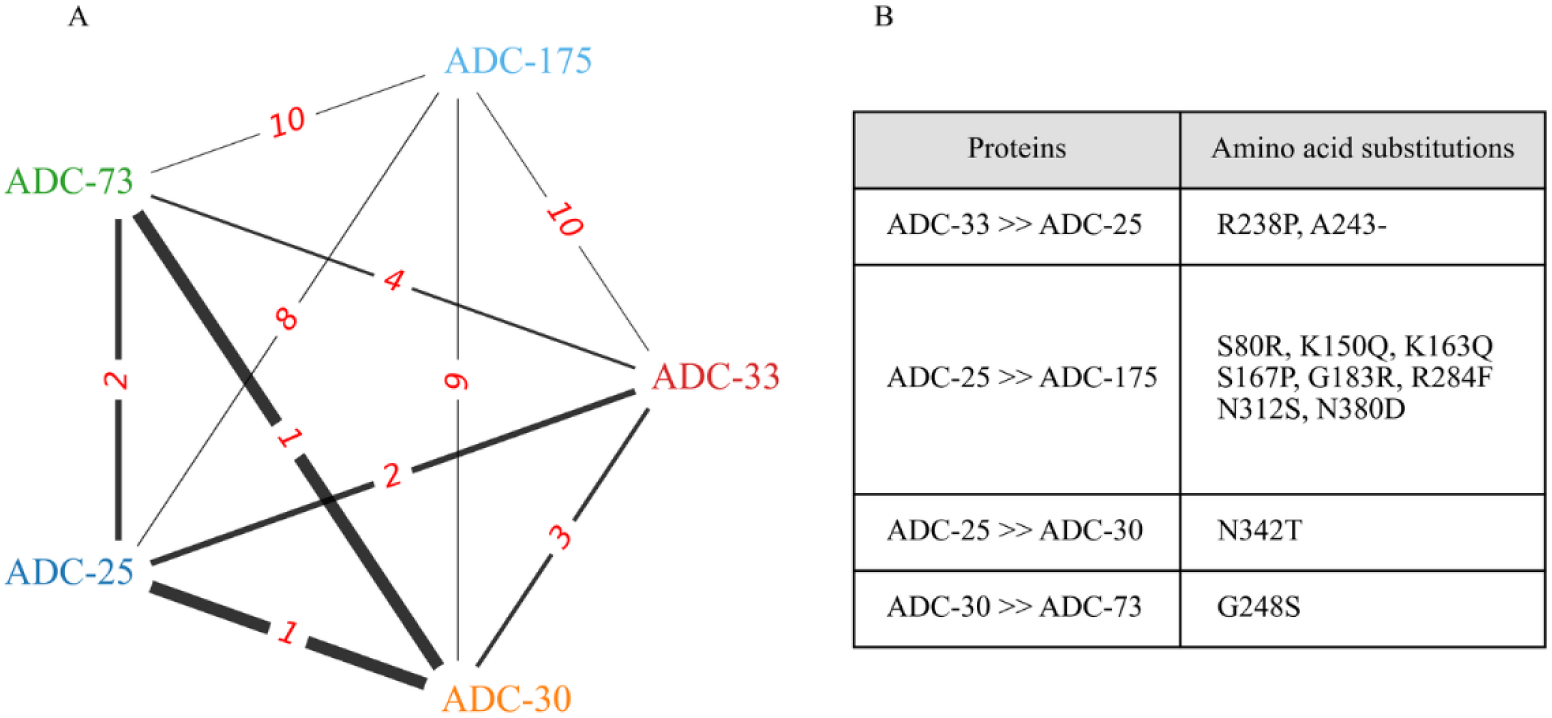
Amino acid differences between major ADC protein variants. (A) Network diagram showing total number of amino acid differences, shown in red, between ADC variants. Line thickness is inversely proportional to the number of mutations between each variant. (B) Table showing specific amino acid differences between selected ADC variants.

The *oxaAb* genes showed less variation, with almost all group 1, 2 and 3 isolates having *oxaAb*(*66*), and almost all group 4 isolates containing *oxaAb*(*82*) (Figure 5). Although these two variants are separated by only a single SNP, this results in an L167V mutation that substantially increases carbapenemase activity through lowered Km (21). However, since group 4 isolates often contain *oxa23*, it is unclear what further advantage this offers. Nonetheless, this confirms that group 4 is distinct from groups 1, 2, and 3.

### Group 4 isolates have few ARGs but many ISs

To further characterise the IC2 groups, ARG, VF, and IS content across chromosomes of isolates from each group were analysed. In line with previous results, group 2 isolates had the most ARGs, followed by group 3, group 1, and finally group 4 (Figure 8A). Further analysis showed that, in addition to the ARGs shown in Figure 4, group 4 isolates often lack other ARGs enriched in IC2, such as *bla_TEM-1_*, *tetR*, and *tet(B)*. Conversely, total VF gene content was similar between groups. Although there was a statistical difference in the distribution of total VFs within chromosomes, Cliff’s delta showed small effect sizes in most cases (Figure 8B), and the high interquartile ranges (IQRs) make it difficult to draw comparisons. Therefore, although ARG content is distinct between different groups, the difference in VF gene content is negligible between groups.

**Figure 8.**
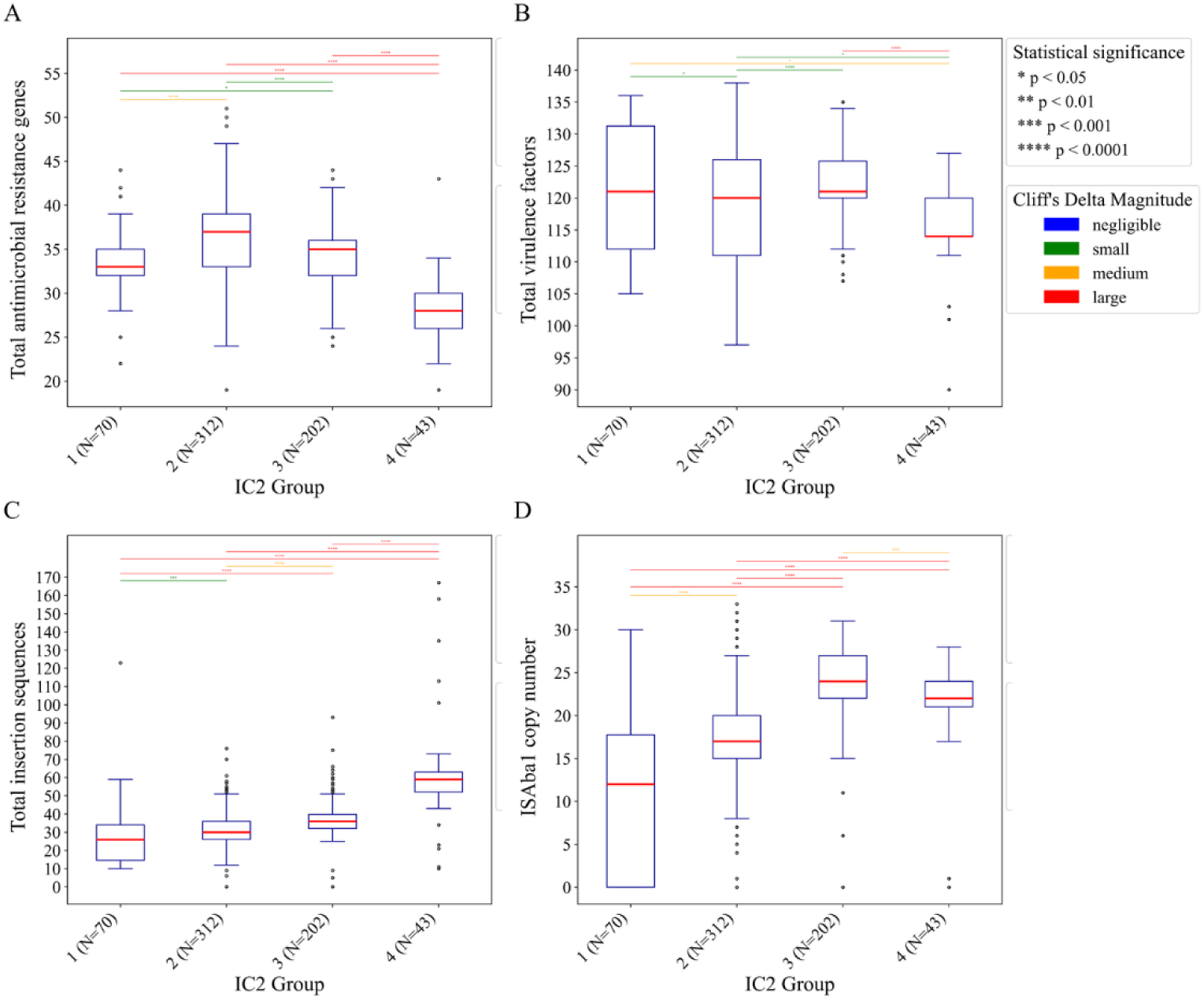
Boxplots of total ARGs, VFs, ISs, and IS*Aba1* copies within chromosomes of IC2 group isolates. Total number of chromosomally encoded ARGs, VFs, ISs, and ISAba1 are shown in (A), (B), (C), and (D) respectively. Medians are shown with red lines. Outliers (±1.5×IQR) are shown as open circles. Statistical significance (*p*<0.05) was determined with the Mann-Whitney U test with Holm-Bonferroni correction applied. Cliff’s delta was used to determine the effect size.

Group 3 chromosomes were found to have the second-highest number of ISs, followed by group 2 and group 1 (Figure 8C). However, group 4 chromosomes had the highest number of ISs (Figure 8C). Further, while group 3 did have the highest median IS*Aba1* copy number, group 4 still had the second highest, followed by group 2, and finally group 1 (Figure 8D). Further analysis showed that the high IS content in group 4 is driven mainly by IS*Aba13*, IS*Aba17*, and IS*Aba125*. The latter of these was commonly associated with the ARG *aph(3’)-VIa* (also called *aphA6*), which in turn was mainly found in isolates dating from after 2022. IS*Aba125* contains an even stronger promoter than IS*Aba1* (22) and likely drives high expression of this ARG, potentially compensating for a lack of *armA* within group 4. Additionally, a median of 56.5% and 64% of IS*Aba13* and IS*Aba17* respectively were found within the K-locus or OC-locus. Therefore, we speculate that the group 4 isolates may have altered capsule and biofilm properties that alter their antimicrobial sensitivity phenotype, which may compensate for a relative lack of ARGs.

## Discussion

While IC2 is the most successful lineage of *A. baumannii*, the genomic factors behind its success and the evolution of its chromosome have remained poorly studied. We leveraged a set of historical *A. baumannii* isolates to identify a set of ARGs that we propose contributed to the success of *A. baumannii* IC2 over IC1. Further, we identify clades within IC2 which share common ARG presence/absence patterns, enabling us to split IC2 into four distinct subgroups.

Analyses of a number of aspects of IC2 suggest that three of the groups identified emerged in sequence over time, with corresponding gain or loss of a subset of ARGs. By collection date alone, the oldest IC2 isolates in our study are found in group 1, and therefore represent isolates that are closely related to the common ancestor with IC1. Consistent with this, group 1 isolates contain ARGs which are also enriched in IC1 (*aadA*, *sul1*, and *qacEΔ1*). Group 2 seems to have expanded around 2005 with the acquisition of a large set of ARGs, many of which are associated with AbGRI3, which first appears in 2005 in this dataset. This is consistent with the proposition that AbGRI3 was originally acquired by IC2 sometime in the early 2000s (15). Our data shows that the rapid expansion of group 2 around 2005 coincides with IC2 overtaking IC1 to become the dominant *A. baumannii* lineage, in line with other reports (18). Therefore, it is probable that the acquisition of AbGRI3 and *oxa23* contributed to the rapid expansion and current success of IC2 over IC1. The rapidly expanding clade 2.5.6 reported by Li *et al*. (18) is equivalent to group 3, which our data also confirms emerged in East Asia around 2005 and is expanding quickly in the USA and China. We found that group 3 began rapidly expanding around 2015 and has lost a subset of ARGs through genomic streamlining in AbGRI3. While some of these lost ARGs have been functionally superseded by other retained ARGs (e.g. *armA* provides broad aminoglycoside resistance), others provided resistance to antimicrobials which are not typically used against *A. baumannii* (*catB8*, *sul1*). Therefore, we propose that the loss of these genes provides a net fitness benefit to group 3 isolates. The proposed transition from group 1 to group 2, then group 3, is also reflected in increasing IS*Aba1* copy number. This aligns with the proposal that the acquisition of IS*Aba1*, and the genomic plasticity that this then engenders, was a major step in the evolution of *A. baumannii* as a pathogen (23,24). Overall, groups 1, 2, and 3 reflect incremental adaptation of the *A. baumannii* IC2 chromosome to AMR selection over time.

In our study, we have identified group 4 as being quite distinct from the other three IC2 groups detected. Group 4 is not described by Li *et al*. (18) as a separate clade, likely due to the majority of these genomes not being present in their data. While the first isolate from group 4 was collected in 2007, *bla_ADC_*, and *oxaAb* results suggest it may have split from other IC2 isolates earlier than this, with the phylogeny suggesting that it is the oldest group and most closely related to the common ancestor with IC1. Further, the subset of ARGs commonly found in this group are *aadA*, *sul1*, and *qacEΔ1*, which suggests these isolates are derived from an early form of IC2 as discussed above. Notably, many isolates from this group contain *oxa23*. Despite lacking many ARGs which are enriched specifically in IC2, the chromosomes of group 4 isolates contain numerous copies of IS*Aba1*, IS*Aba125*, IS*Aba13*, and IS*Aba17*. Colocalization between *oxa23*/IS*Aba1* and *aph(3’)-Via*/IS*Aba125* has been described in the literature conferring resistance to carbapenems and amikacin respectively (25,26). Amikacin in particular is an aminoglycoside often used to treat infections by AMR Gram-negative bacteria, including *A. baumannii* (27,28). Additional copies of IS*Aba13* and IS*Aba17* inserted into the K-locus and OC-locus may provide further resistance through disruptions of genes involved in capsule and biofilm biosynthesis. IS-mediated disruption of genes involved in these pathways, especially by IS*Aba13*, has been previously described, in some cases with corresponding increases in virulence and/or antimicrobial resistance (29–31). Alterations in the capsule and biofilm formation may reduce the need for ARGs against specific antimicrobials, compensating for the general lack of ARGs in these isolates. These mechanisms, some of which are rarely seen in IC2 isolates outside of group 4, may have enabled group 4 to persist at low levels while its closest counterpart, group 1, became comparatively rare. There has been a notable increase in the isolation and sequencing of group 4 isolates since 2023, mainly driven by pathogen surveillance programs in the USA. While it is hard to determine if this is due to increased sampling, we note similarities to the pattern of emergence in group 2 and group 3, and the periodicity at which new forms of IC2 emerge. Therefore, group 4 may be in the early stages of expansion.

We note that historical isolates sequenced and assembled here show a bias towards the UK and Europe. This resulted in a notable lack of isolates from Asia and the Middle East, despite these regions having *A. baumannii* outbreaks in the early 2000s (32). This bias may have offset the detected date when a gene was first acquired by a few years if the gene was first acquired in these regions. For instance, the first reported occurrence of *armA* in *A. baumannii* was in Korea in 2003 (33), while the first occurrence in our dataset was in China in 2005. Nonetheless, these historical chromosomes provide significant insight into the evolution of the *A. baumannii* IC2 chromosome during its emergence which would not be possible with pre-existing data. Additionally, plasmid sequences in publicly available genomes were analysed for ARG/VF content. Almost all plasmids contained no VFs and either one or no ARGs (data not shown) in line with previous studies (6–8). Hence, plasmids were excluded from this study.

In conclusion, we leveraged a set of historical *A. baumannii* isolates to identify patterns of ARG gain and loss which may have contributed to the success of IC2 over other *A. baumannii* lineages. Further, we find that IC2 can be split into at least four distinct groups, with matching ARG presence/absence patterns. While group 1 represents an early form of IC2, we propose that group 2 became highly successful in part due to the acquisition of *oxa23* and the formation of AbGRI3. Group 3 contains a streamlined form of AbGRI3, which we propose has enabled it to rapidly expand within the last decade. Group 4, meanwhile, is distinct from other groups with a parallel evolutionary history that may be in the early stages of expansion.

## Methods

### Historical isolates sequencing

Following overnight growth at 37 °C in Miller’s lysogeny broth, genomic DNA was extracted using the Quick-DNA Magbead Plus Kit (Zymo Research). DNA was purified further using Ampure XP beads (Beckman Coulter). DNA libraries were prepared using the Rapid Barcoding Kit 96 V14 from Oxford Nanopore Technologies (ONT). Sequencing was done with an R10.4.1 PromethION Flow Cell using a PromethION 2 Solo. Base calling and adaptor trimming was performed using Dorado v0.7.2 with the dna_r10.4.1_e8.2_400bps_sup@v5.0.0 model (ONT, Oxford UK). Reads under 1 Kbp or below Q10 were filtered out using Filtlong v0.2.1 (34). Samples with predicted coverage <30X were removed, and samples with a predicted coverage >80X were downsampled using Filtlong. Read length was prioritised over read quality by setting length weight to 10 and leaving quality weight at default of 1.

### Historical chromosome assembly and quality control

All computational analysis was performed on a high-performance computing (HPC) cluster running Rocky Linux v8.10. Reads were assembled using Flye v2.9.5-b1801 (35) with the nano raw model. Target coverage was set to 50X with an estimated genome size of 3.9 Mbp. A single round of polishing was performed with the in-built polisher. Next, one round of Racon v1.5.0 (36) and one round of Medaka v2.0.1 (Oxford Nanopore Technologies, 2018) polishing was applied to all assemblies with Minimap v2.28-r1209 used as the aligner. Finally, ReCycler v0.1.0 (37) was used to circularise and start all contigs at the *dnaA* gene, with a small subset of 20 contigs where this process failed being kept.

BUSCO scores, which assess the completeness of the annotations by identifying a set of highly conserved single-copy orthologs, were obtained for the longest contig in each assembly using BUSCO v5.8.3 (38). Kraken v2.1.4 (39) was used to classify each contig. Only contigs classified as *A. baumannii*, >3.5 Mbp and <4.5 Mbp in length, having GC content >36% and <43%, containing at least one origin as detected by ReCycler, and with BUSCO scores >95% were considered whole chromosomes. ANI, which evaluates the genetic relatedness of two genomes, against the *A. baumannii* type strain (ATCC 19606, GenBank accession no. CP045110.1) was checked using FastANI v1.34 (40), with all isolates having >95% ANI.

### Publicly available genome retrieval and metadata collation

Publicly available *A. baumannii* genomes marked as ‘chromosome’ or ‘complete genome’ were retrieved using the National Centre for Biotechnology Information (NCBI) datasets utility (41) (Accessed 24/05/2025). Genomes marked as atypical or suppressed were removed. For each genome, the longest contig was extracted, and any contigs shorter than 3 Mbp, longer than 5 Mbp, or not containing a *dnaA* gene were removed. Metadata was processed to obtain the collection year and country. Where a range of years was given, the average was taken. This resulted in a total of 1,055 publicly available chromosomes which were used to supplement the 226 historical chromosomes.

### Chromosome annotation

Chromosomes were annotated with bakta v1.11 (42) using the light database. ARGs were annotated with the RGI v6.0.4 which queries the CARD (43). VFs were annotated using ABRicate v1.0.1 (44) which queries the virulence factor database (VFDB) (45). ISs were annotated using digIS v1.2 (46). Integron features were annotated using IntegronFinder v2.0.6 with genome topology set to circular and local max set to true for increased sensitivity (47).

### Core genome phylogeny

Panaroo v1.5.2 (48) was run on GFF3 files output by Bakta to create a core genome alignment with MAFFT (49) as the aligner and core genome threshold of 1. IQtree v2.4.0 (50) was run on this alignment with the substitution model determined using ModelFinder (51). The core genome phylogeny was rooted with an *Acinetobacter nosocomialis* outgroup (GenBank accession no. CP157432.1).

### MLST and IC assignment

*In silico* MLST was performed on all genomes using FastMLST v0.0.16 (52) using the Pasteur scheme. Clonal complexes were constructed at the single locus variant (SLV) level using the goeBURST algorithm (53) implemented in PHYLOViZ (54). The founder ST, and other associated STs, for each IC was obtained from previously published data (4,55,56). Isolates which were both assigned to the same clonal complex and monophyletic group within the core genome phylogeny as an isolate of a founder ST were assigned to the corresponding IC.

### Recombination-free IC2 phylogeny

Whole genome alignment of chromosome sequences was performed using ska2 v0.4.0 (57) against a recent ST2 isolate (GenBank accession no. CP181411.1). Gubbins v3.4 (58) was run with the first tree builder set to FastTree and subsequent tree builder set to IQtree. First model was set to JC, with subsequent models being computed using IQtree with RAxML used as the model fitter. Minimum SNPs was set to 2 for aggressive removal of recombination, and marginal state reconstruction was used. To obtain support values, 1,000 bootstrap replicates were performed. The ATCC 19606 genome was used as an outgroup, and was pruned from the final tree using newick utilities v1.6 (59). Visualisation was performed using TreeViewer (60).

### Temporal and other statistical analysis

Gene enrichment/depletion was determined based on ARG annotations produced by RGI with Fisher’s exact test using Scipy v1.13.1 (61). Root-to-tip analysis was performed using BactDating v1.1.3 (62) in R v4.3.1. Nodes without dates were pruned before analysis using ape v5.8-1 (63). Mann-Whitney U test was performed to determine if distributions of total features (ARGs, VFs, ISs, ISA*ba1*) were significantly different between groups. To quantify the size of this difference, effect size was calculated with Cliff’s delta (64). All *p* values were adjusted for multiple testing using the Holm-Bonferroni correction with statsmodels v0.14.4 (65).

### ADC variant analysis

Amino acid sequences for ADC-175, ADC-25, ADC-30, ADC-73, and ADC-33 were obtained from the CARD (CARD accession no. WP_001211208.1, ABK34773.1, AEL30572.1, ALA14808.1, WP_001211220.1 respectively). Alignment was performed using MUSCLE v5.3. (66). Network analysis was performed using NetworkX v3.1 (67).

## Author Contributions

MN: conceptualization, methodology, formal analysis, investigation, data curation, writing – original draft preparation, visualization; FG: investigation, data curation, writing – review and editing; NA: investigation, data curation, writing – review and editing; GCL: conceptualization, methodology, writing – review and editing, supervision; SCR: conceptualization, methodology, resources, data curation, writing – review and editing, supervision; LPH: conceptualization, methodology, resources, data curation, writing – review and editing, supervision, project administration, funding acquisition; BAE: conceptualization, methodology, resources, data curation, writing – review and editing, supervision, project administration, funding acquisition.

